# Functional decoupling of emotion coping network subserves automatic emotion regulation by implementation intention

**DOI:** 10.1101/336800

**Authors:** Shengdong Chen, Nanxiang Ding, Bharat Biswal, Zhihao Li, Shaozheng Qin, Chao Xie, Changming Chen, Jiajin Yuan

## Abstract

Automatic emotion regulation (AER) aims at modifying emotional impact effortlessly. However, the effortless account and the neural mechanisms of AER are both undetermined. For this purpose, we collected functional neuroimages (fMRI) in thirty-one participants who attended to neutral and disgust pictures in three conditions: Watching, Goal Intention (GI) and Reappraisal by Implementation Intention (RII). RII decreased negative feelings and bilateral amygdala activity without eliciting cognitive efforts, evidenced by the reduced effort rating and less prefrontal engagement during RII compared to Watching and GI. These regulation effects should not be explained by emotional habituation, as Experiment 2 (n=40) observed no habituation to stimulus repetitions. Task-based network analysis showed similar functional connectivity (FC) of ventral anterior cingulate cortex to left insula and right precuneus during RII and GI conditions, both involving goal setup. Furthermore, RII relative to GI exhibited weaker FC in brain networks subserving effortful control (e.g. inferior-superior parietal FC), memory retrieval (e.g. inferior-middle temporal and lingual-putamen FCs), aversive anticipation and motor planning (e.g. Paracentral-superior temporal gyrus, putamen-operculum FCs). The FC strength of putamen to operculum/lingual, and paracentral to STG positively predicts regulatory difficulty. These results suggest that the setup of implementation intention automatizes emotion regulation, by reducing online mobilization of neural systems underlying the stream of emotion coping.

## Introduction

Emotion regulation, a process to modulate any components of emotional activity (Gross, 1998), plays a vital role in the maintainance of one’s health and social functioning. Though it is suggested that emotion regulation can be realized by either effortful or automatic process (Gross, 1998, 2014), most studies to date focus on the voluntary forms of emotion regulation that is resource-demanding, such as intentional reappraisal (Goldin, McRae, Ramel, & Gross, 2008; Ochsner, Bunge, Gross, & Gabrieli, 2002). For instance, a number of studies observed that intentional reappraisal decreased emotional experience and emotion-related subcortical activation (e.g. amygdala), at the cost of increasing control-related prefrontal activation (Buhle et al., 2014; Goldin et al., 2008; Ochsner et al., 2002). This cognitive cost, in certain cases, may lead to the failure of emotion regulation. For example, individuals with generalized anxiety disorder or major depressive disorder are characterized by deficits in prefrontal cognitive control function (Joormann & Gotlib, 2010; Rogers et al., 2004), which consequently leads to disinhibition of negative emotion (Disner, Beevers, Haigh, & Beck, 2011; Joormann & Gotlib, 2010).

Recently, an increasing number of behavioral and electrophysiological studies have examined automatic form of emotion regulation (Chen, Deng, et al., 2017; Gomez, Scholz, & Danuser, 2015; Williams, Bargh, Nocera, & Gray, 2009; Yuan, et al., 2015a and 2015b), using method like implementation intention (Gallo, McCulloch, & Gollwitzer, 2012; Gomez et al., 2015). It has been suggested that the antecedent buildup of implementation intentions leads to automatic attainment of emotion-regulatory goals, without mobilization of intentional regulatory efforts upon emotional stimulation (Eder, 2011; Gallo, et al., 2009 and 2012). In an early study, Gollwitzer (1999) differentiated two kinds of intentions: goal intention and implementation intention. Goal intention (GI) defines a desired end state and has a format of “I want to attain Z!”. However, people often failed to achieve emotion-regulatory goals when instructed only to set a goal intention (e.g., “I will not get upset!”) (Gallo & Gollwitzer, 2007a; Gomez et al., 2015). Implementation intention, by contrast, is formed to realize goal intention by specifying when, where, and how goal-directed behaviors should be initiated in the form of “if-then” plan (e.g. “If loss sign is encountered, then I will keep calm!”). Thus, implementation intention links a goal-relevant situational cue (e.g. loss sign) with a goal-directed behavior (e.g. “keep calm”), which reduces the intention-behavior gap between intended outcome and actual goal attainment (Gollwitzer & Sheeran, 2006). The effects of implementation intention are explained by that the mental link created between critical cue and behavior turns the effortful and top-down control of goal-directed responses, into an automated, effortless and bottom-up stimulus control (Gollwitzer, 1999; Webb & Sheeran, 2008).

Gallo and Gollwitzer (2007a) first reported that forming implementation intention for the control of spider phobia reduced the subjects’ fearful experience for spider-related stimuli during a cognitive demanding task. Additionally, an Event-Related Potentials (ERP) study by Gallo et al (2009) reported that forming an implementation intention reduced occipital P1 amplitudes for threatening stimuli compared to GI or watching condition. Recently, Gallo et al. (Gallo et al., 2012) found that a reappraisal-based implementation intention allowed participants to rate disgusting pictures as less unpleasant than participants in the watching or GI groups. However, these behavioral and electrophysiological studies of implementation intention did not use objective indexes to assess cognitive costs between regulation and no-regulation conditions, thus unable to verify the effortless or automatic account. Though Gallo and colleagues (Gallo & Gollwitzer, 2007a; Gallo et al., 2009 and 2012) have collected data of subjective efforts and experienced difficulty during goal and implementation intention, these data were not collected during the control condition. In brief, these findings suggest that implementation intentions can regulate emotional responses effectively, but leave it unknown whether implementation intention or goal intention enhanced cognitive efforts compared to the baseline, watching condition.

Further, what these studies measured are self-reported or electrophysiological variables. Few studies to date have examined neural mechanisms of automatic emotion regulation by implementation intention. To the best of our knowledge, there is only one fMRI study that involved implementation intention and emotion regulation (Hallam et al., 2015). This study observed reduced self-reported affect and amygdala activity during implementation intention compared to goal intention. However, the reappraisal strategy used in this study was performed effortfully rather than effortlessly, as participants were reminded to reappraise the picture meanings every time a reappraisal cue was presented which, as a result, elicited prominent dorsolateral prefrontal activation subserving voluntary inhibition (Hallam et al., 2015). Therefore, it is highly necessary to examine the effectiveness of automatic emotion regulation by implementation intention, in terms of both emotional impact and cognitive costs. Behavioral and neuroimaging measures are both used to test this hypothesis. Moreover, if the automaticity of emotion regulation is verified during RII, the neural mechanisms underlying the automatic emotion regulation needs to be further investigated. We performed two experiments, one functional MRI and the other behavioral, to answer the above questions.

## Experiment 1

### PURPOSE AND RATIONALE

This experiment aims to examine whether Reappraisal by Implementation Intention (RII) can regulate emotional consequences automatically, without increasing cognitive costs. Cognitive reappraisal, which requires reformulating the meaning of the emotional situation, was chosen as the target strategy to be planted into implementation intention, because the regulatory efficacy and cognitive costs of this strategy are both established (Goldin et al., 2008; Gross, 2002; Richards & Gross, 2000). Disgust was chosen as the target emotion, for it has proven to elicit robust neural activation in both cognitive-control and emotion-generative regions (e.g. amygdala; Goldin et al., 2008; Schienle, et al., 2005; Wicker et al., 2003).

In addition, empirical studies and recent meta-analysis suggest that amygdala and insula are central regions underlying the generation of disgust (Goldin et al., 2008; Schienle et al., 2005; Wicker et al., 2003), and that prefrontal regions like dorsolateral prefrontal cortex (dlPFC) and dorsal anterior cingulate cortex (dACC) are involved in voluntary reappraisal and top-down cognitive control (Buhle et al., 2014; Goldin et al., 2008; Kalisch, 2009; McRae et al., 2010, 2012; Ochsner et al., 2002). Therefore, neural activity in these two emotion-generative regions (amygdala and insula) and two cognitive control-related regions (dlPFC and dACC) provide objective indexes of emotional responses and cognitive costs, respectively. If RII regulates emotion automatically, it should reduce the activity in emotion-generative regions without increasing the activity of the cognitive control regions compared to GI or control condition. We also conducted a voxel-wise whole-brain analysis to test whether other regions were involved in RII in addition to the regions of interest. If the automatic/ effortless regulation is confirmed, a task-related network analysis would be performed to explore the neural mechanisms subserving the automatic emotion regulation by RII.

## METHODS AND MATERIALS

### Participants

Thirty-one healthy, right-handed college students from the Southwest University in China with normal or corrected-to-normal vision participated in this study. We determined the sample size based on a power analysis using G-power software (Faul et al., 2009). We specified moderate effect size (0.25), 0.8 power, and a moderate correlation (0.5) among the repeated measurements, which yielded a recommended sample size of 28. We collected 31 participants’ fMRI data to avoid the possibility that some of data cannot be used because of head movement or other potential reasons. Participants’ mean age was 21.34 years, ranged from 19 to 25. All participants reported no history of neurological or psychiatric disorders. Written informed consent was obtained before the experiment in accordance with the Institutional Review Board of the Southwest University and the latest revision of the Declaration of Helsinki. Data of 5 participants were excluded due to excessive head movement (larger than 3 mm) during fMRI scanning, resulting in the inclusion of 26 participants (16 males and 10 females).

### Stimuli

The stimulus material consisted of two categories with 90 pictures in total: 45 disgust and 45 neutral pictures, taken from the International Affective Picture System (IAPS) (Lang, Bradley, & Cuthbert, 1999) and the Chinese Affective Picture System (CAPS) (Lu, Hui, & Yu-Xia, 2005). The valence and arousal scores of each picture were assessed by 30 independent raters, independent to the experiment sample. The disgusting pictures showed bloody burn victims and mutilated bodies. Within the bi-dimensional model of valence and arousal, such contents are rated as negative and high-arousal, while neutral pictures had medium standard emotional valence and low arousal ratings. The pictures were presented in a randomized order and the raters were asked to rate to what degree they felt sadness, fear, joy, disgust, and anger on scales ranging from 1 (little) to 7 (very). Results revealed a significant main effect for the unpleasant pictures, (F(4,40)=1088.96, p<0.001, η^2^=0.96), Post hoc bonferroni tests showed that disgust was the most prevalent emotion (M = 5.74) in comparison with fear (M = 4.22, p<0.001), sadness (M= 3.90, p<0.001), anger (M =2.85, p<0.001) and joy (M = 1.41, p<0.001). These findings suggest that the unpleasant pictures elicit disgust effectively.

### Design and Procedure

Prior to fMRI scanning, participants were trained to be familiar with the experimental task through viewing 15 independent neutral pictures. Participants were told to estimate their positive and negative emotional intensity respectively, after the presentation of 3 consecutive pictures using a 5 point-scale ranging from 0 (not at all) to 4 (very), “How positive or negative did you feel?” After viewing all the pictures, participants received a questionnaire that assessed the consumption of cognitive resources: “How much did you try to cope with negative feelings?” and “How difficult was it to cope with negative feeling?” The two items were also accompanied by a 5 point-scale ranging from 0 (not at all) to 4 (very).

The present study used a 3×2 factorial design with the regulation condition (watching, GI and RII) and type of pictures (neutral/negative) as repeated factors. The fMRI task consisted of three ordinal runs corresponding to three conditions: passive watching, goal-directed intention and implementation intention. Each run consisted of 10 blocks (5 neutral and 5 negative blocks) that matched in valence and arousal, and each block consisted of 3 consecutive pictures (2s each) of the same valence. Both neutral and negative pictures across the three conditions were not significantly different in valence (F_neutral_(2, 43)=0.03, p_neutral_=0.97, η^2^_neutral_=0.01; F_negative_ (2, 43) =0.015, ρ_negative_=0.99, η^2^_negative_=0.01) and arousal ( F_neutral_ (2, 43) =0.19, ρ_neutral_=0.84, η^2^_neutral_=0.02; F_negative_ (2, 43) =0.69, ρ_negative_ =0.51, η^2^_negative_=0.03).

In the watching condition (run1), participants were just required to pay close attention to the pictures without further instructions. In the GI condition, participants were additionally asked to form the GI “I will not get disgusted!”. In RII condition, participants were further required to form the following if-then plan: “And if I see blood, then I will take the perspective of a physician!” in addition to the close attention and the formation of GI. Participants received the instructions of RII and reinforced it by rehearsal for one minute. In the formal experiment, in order to remove any additional voluntary regulatory process, participants were only required to attend to the pictures without any further task. Each of the 10 blocks was presented for 6s in a random order following a fixation of 6 to 10s (average 8s). Following 10 block, two scales for the assessment of positive and negative emotion intensity appeared on the screen for 4s each. At the end of each run, two scales assessing effort in emotion control were also presented for 4s each.

### Imaging Data Acquisition and Preprocessing

Whole brain T2*-weighted echo planar imaging was performed with a Siemens Trio 3.0 Tesla (Magnetom Trio, Siemens, Erlangen, Germany) scanner with a gradient echo planar imaging sequence (32 axial slices, TR/TE==2 s/30 ms, FA = 90°, matrix size=64×64, FOV =220×220 mm^2^, voxel size= 3.4×3.4×3 mm^3^, 386 volume measures). High-resolution structural images were acquired for registration purposes using a T1-weighted magnetization-prepared rapid gradient-echo (MP-RAGE) sequence (TR/TE = 1900 ms/2.52 ms, FA = 9°, FOV = 256 × 256 mm^2^; slices = 176; thickness = 1.0 mm; voxel size = 1 × 1 × 1 mm^3^).

MRI data analysis was performed using Statistical Parametric Mapping (SPM8; www.fil.ion.ucl.ac.uk/spm) and custom-written programs in Matlab. Functional images were rigid-body motion corrected and the mean image was coregistered to each participant’s anatomical MR image. Then, the coregistration parameters were used to register all aligned functional scans to the T1. Subsequently, images were transformed into the common MNI space by warping individual’s T1 images and resampled into 3 mm isotropic voxels. Finally, normalized images were spatially smoothed with a Gaussian kernel of 8-mm full width at half maximum. Head movement estimates derived from the realignment step were included as nuisance regressors in subsequent general linear modeling (GLM) to help diminishing the impact of movement-related effects.

### Imaging Data Analysis

For each participant, the voxel-wise whole brain GLM included 6 regressors of interest (negative and neutral pictures in three scans of watching, GI and RII). At the group level, a general linear contrast of watching-negative versus watching-neutral was applied to detect brain activation associated with disgust responses. Based on Random Field Theory, T-statistics for each voxel were thresholded at p < 0.01 and an extent of 10 voxels for multiple comparisons across whole brain with a family wise error rate (FWE).

Region of interest (ROI) analyses were next performed to test whether RII could decrease activation related to disgust responses in typical emotion-generative regions. The emotion-generative regions (bilateral amygdala and bilateral insula) were defined by the respective anatomical masks of the AAL atlas (Tzourio-Mazoyer et al., 2002) by WFU_PickAtlas toolbox (Maldjian, Laurienti, Kraft, & Burdette, 2003). And to examine whether RII increase activation in cognitive-control related regions, three ROIs in bilateral dlPFC and dACC were further defined as the Brodmann areas 9 and 46 combined (left and right), as well as a 10 mm radius sphere at Talairach coordinates (x = 0, y = 12, z = 42) (Shackman et al., 2011), respectively. For each of these ROIs, mean Percent Signal Change (PSC) of each individual was extracted using Marsbar toolbox (Brett, Anton, Valabregue, & Poline, 2002). Then, the emotional intensity of negative affect during each condition was represented as the average contrast values (negative > neutral) for the four emotion-generative regions and three cognitive control-related regions.

### Estimation of task-related FC

Using the CONN toolbox (version16, www.nitrc.org/projects/conn) in MATLAB, we estimated the task-related FC between each pair of brain regions in a network of 229 spherical (radius=3mm) regions. Of these regions, 227 ROIs were selected from 264 coordinates reported by Power et al. (2011). Those coordinates are the centers of putative functional areas (and subcortical and cerebellar nuclei), defined by multiple task fMRI meta-analyses (Dosenbach et al., 2010) and by a resting state FC MRI parcellation technique (Cohen et al., 2008). These 227 ROIs have also been assigned to 10 well established functional networks, comprising low-level input and output networks (visual, auditory and sensorimotor networks), subcortical nodes, the default mode network (DMN), ventral and dorsal attention networks (VAN and DAN), and cognitive control networks (Frontal-Parietal network, FPN; cingulo-opercular network, CON, salience network, SN) (Cole et al., 2013; Mohr et al., 2016). The two ROIs added in the connectivity analysis were bilateral amygdala defined via the contrast of watching-negative > watching-neutral (**Table 1**). They were specifically included for their central role in the processing of disgust. Details of these ROIs (coordinates and labels) are provided in **S1 Data**.

**Table 1.** Group Activations for Contrast Watch-negative Versus Watch-Neutral

| Brain Regions | Brodmann | x | y | z | Voxels | t-Value |
| --- | --- | --- | --- | --- | --- | --- |
| **Watch-negative>Watch-Neutral** |  |  |  |  |  |  |
| Frontal Lobes |  |  |  |  |  |  |
| L Anterior Cingulum | 32 | -12 | 42 | 6 | 94 | 11.46 |
| R Medial Superior Frontal | 10 | 3 | 63 | 18 | 576 | 10.57 |
| R Inferior Frontal Orbitalis | 47 | 39 | 33 | -18 | 37 | 9.69 |
| L Medial Superior Frontal | 10 | -12 | 63 | 18 | 24 | 9.59 |
| R Insula | 48 | 24 | 15 | -18 | 240 | 9.54 |
| L Middle Cingulum | 23 | -12 | -24 | 36 | 38 | 9.50 |
| R Inferior Frontal Triangle | 48 | 36 | 18 | 18 | 304 | 8.71 |
| L Insula | 48 | -36 | 6 | -6 | 67 | 8.43 |
| L Rectus | 11 | -6 | 39 | -18 | 91 | 5.52 |
| Temporal Lobes |  |  |  |  |  |  |
| R Middle Temporal Lobes | 37 | 48 | -72 | 3 | 948 | 14.63 |
| L Middle Temporal Lobes | 37 | -57 | -57 | 6 | 847 | 11.35 |
| R Fusiform | 19 | 27 | -57 | -12 | 364 | 11.16 |
| Parietal Lobes |  |  |  |  |  |  |
| L Postcentral | 3 | -33 | -39 | 54 | 108 | 8.28 |
| L Inferior Parietal | 3 | -45 | -24 | 39 | 334 | 7.46 |
| Occipital Lobes |  |  |  |  |  |  |
| R Middle Occipital | 37 | 48 | -72 | 3 | 142 | 14.63 |
| L Middle Occipital | 19 | -48 | -78 | 6 | 941 | 12.49 |
| R Lingual | 18 | 15 | -72 | -15 | 128 | 10.74 |
| L Lingual | 19 | -24 | -66 | -9 | 38 | 8.16 |
| Subcortical Regions |  |  |  |  |  |  |
| L Thalamus |  | -27 | -21 | -6 | 33 | 11.53 |
| R Hippocampus |  | 24 | -6 | -18 | 135 | 10.66 |
| R Amygdala |  | 30 | 0 | -21 | 27 | 10.46 |
| L Hippocampus |  | -24 | -9 | 15 | 273 | 10.35 |
| L Amygdala |  | -24 | -6 | -15 | 220 | 10.18 |
| **Watch-Neutral > Watch-negative** |  |  |  |  |  |  |
| No significant clusters of activation |  |  |  |  |  |  |

*Note.* All clusters reached a significance level of $p=0.01$ , FWE corrected and an extent threshold of 10 voxels. For each cluster, x, y, z, MNI coordinates; L, left; R, right.

Regional time series within each of these 229 ROIs were extracted from the preprocessed fMRI data on individual level. The task onset times were modeled, and covariates of no interest (e.g. the realignment confound, white matter and CSF signal) were regressed out using a component-based noise correction method (CompCor) (Behzadi, Restom, Liau, & Liu, 2007). Time series of voxels within 229 ROIs were averaged, and those average time series were correlated with each other. The resulting correlation coefficients were then Fisher z-transformed to normalize their distribution. These values represent the connectivity between the source and target regions during each task condition. The computed ROI-to-ROI connectivity matrices of each participant were finally entered into the second-level group analysis that treated participants as a random variable in a 3-by-2 ANOVA. False positives in this network analysis was controlled by false discovery rate (FDR) of P < 0.05.

## RESULTS

### Manipulation Check of Disgust Induction

To check whether the target emotion disgust was elicited by the disgusting pictures, we contrasted watch-negative versus watch-neutral on both experience and neural indices of negative emotion during the passive watching. Results showed that the watch-negative versus watch-neutral contrast resulted in increased subjective ratings of negative affect (t(25)=28.49, p<0.01) (**Figure 1A**), and increased responses in prefrontal cortex, bilateral temporal and occipital cortex, parietal cortex, and subcortical regions (**Table 1**). As expected, we confirmed that typical emotion-generative regions, including amygdala and insula, were bilaterally activated by this contrast (**Figure 1B**). There were no significant brain responses for watch-neutral versus watch-negative contrast. Both of behavioral and neural findings supported that the target emotion disgust was successfully elicited by the disgusting pictures.

**Figure 1.**
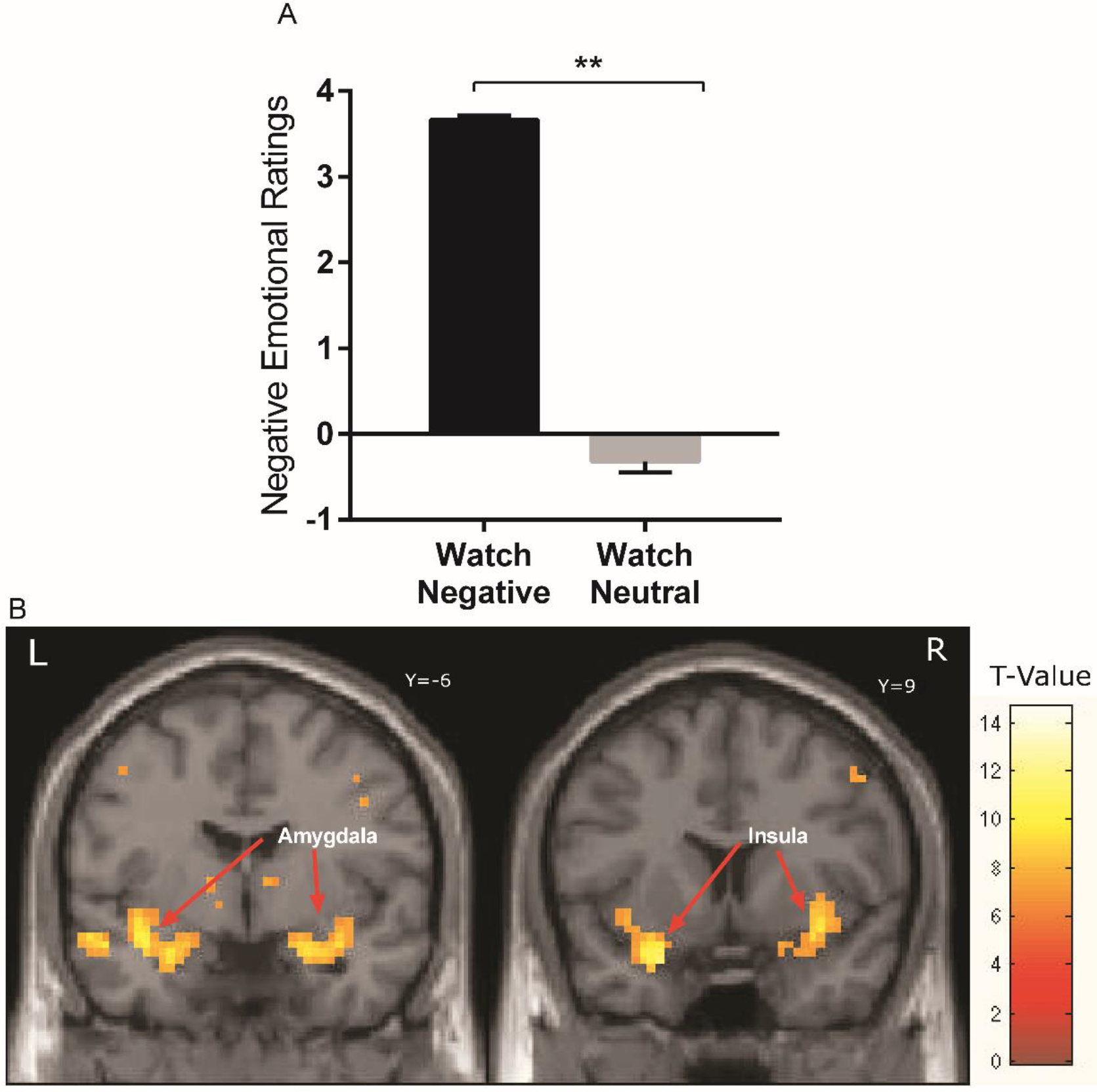
The contrast of watch-negative versus watch-neutral on behavioral and neural indices. (**A**) Negative emotional ratings during passively viewing of disgust pictures (watching condition). Error bars= SE. (**B**) From the one-sample t-test across all 26 participants for the contrast watch-negative versus watch-neutral. The display threshold was p=.01, FWE corrected and an extent of 10 voxels. ** means p≤0.01

### Emotion Regulatory Effects of RII: behavioral and ROI analyses

**Figure 2.**
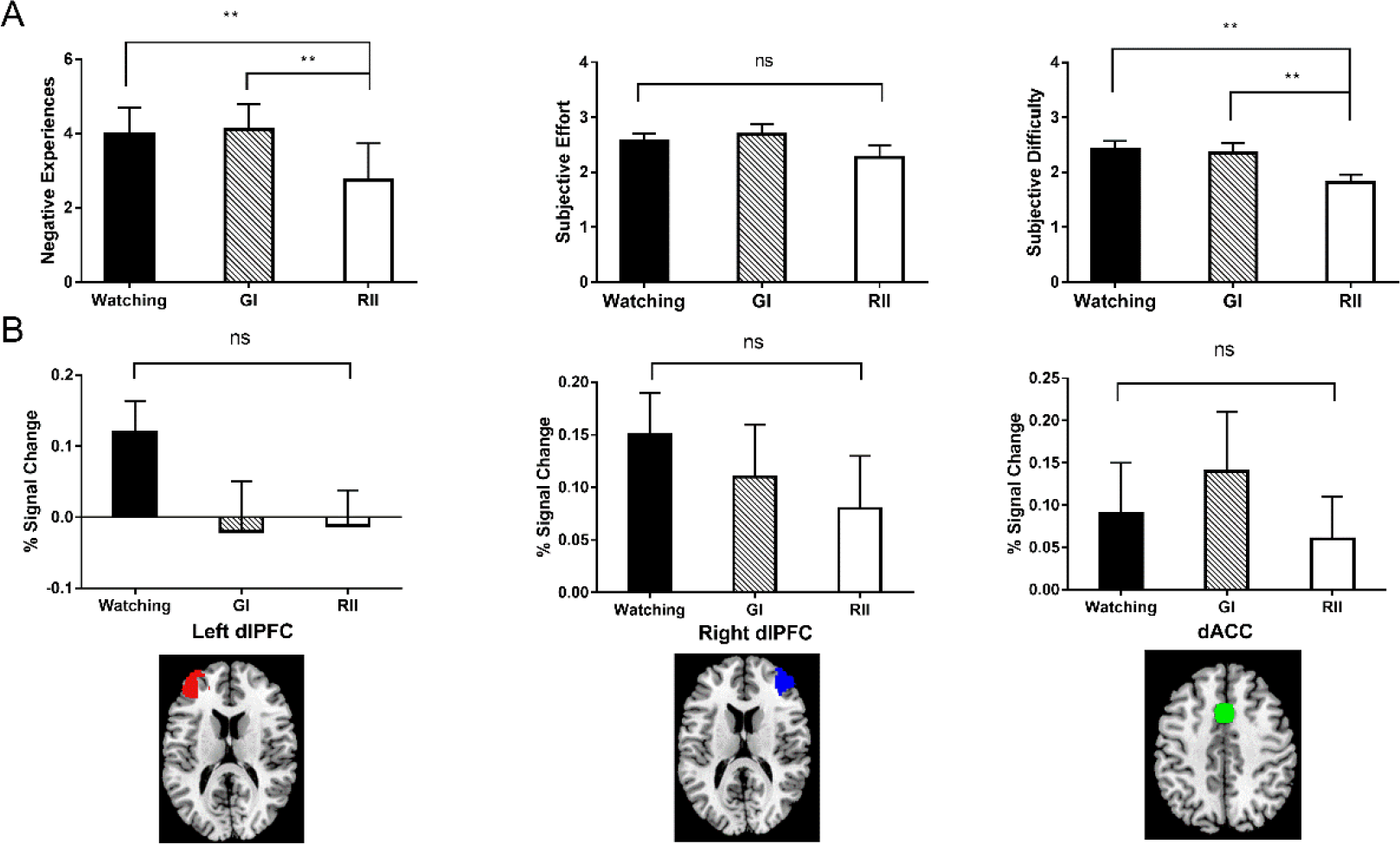
(**A**) The subjective negative experience, the subjective effort and subjective difficulty in negative emotion regulation among the watching, GI and RII conditions. (**B**) Mean percent BOLD signal changes of control-related regions across the three conditions. The cognitive costs during each condition were represented by the PSC during the disgust block minus during the neutral block (baseline). Error bars=SEM, ** means p≤0.01, ns stands for not significant.

**Figure 3.**
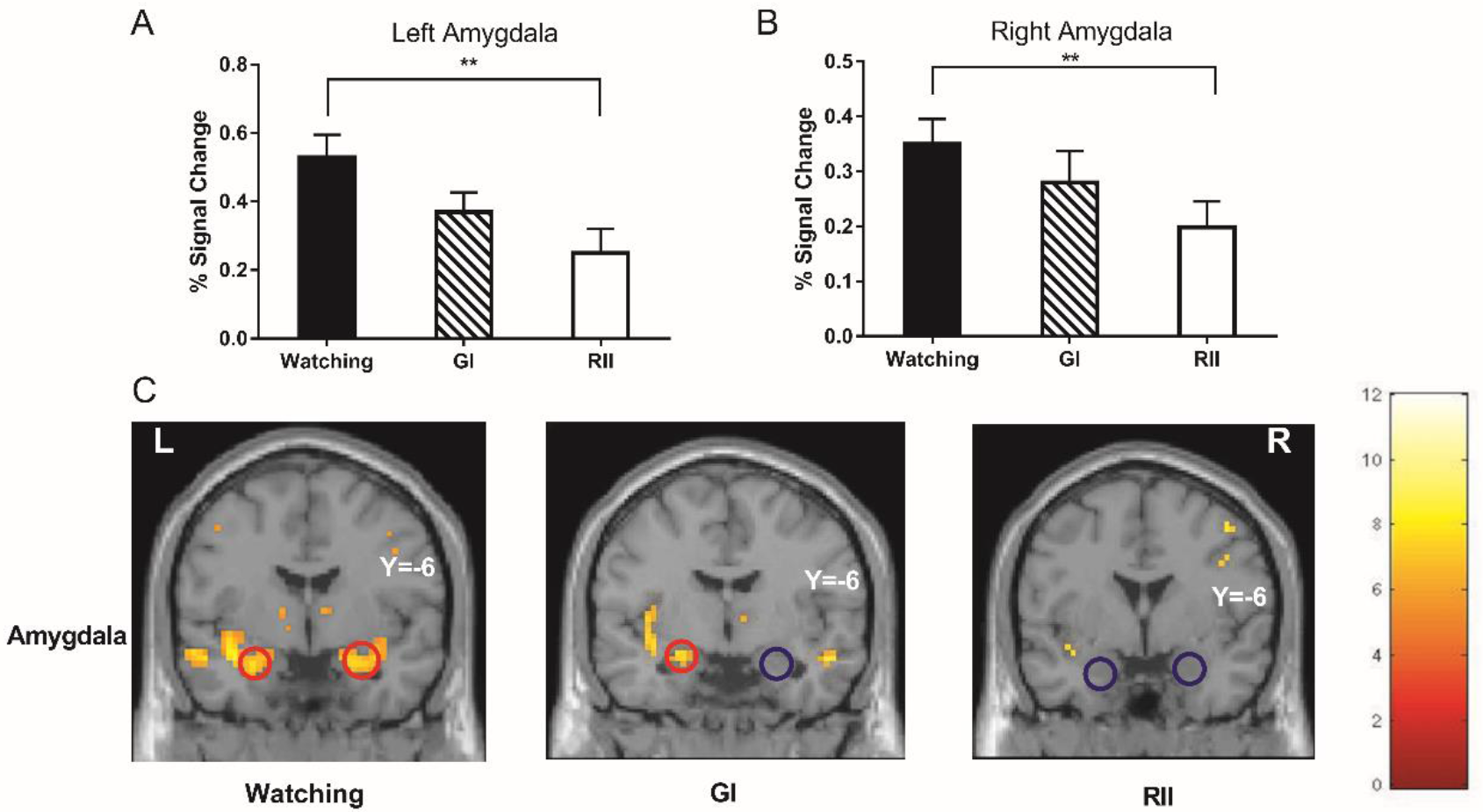
The regulatory effects of RII in the left (**A**) and right amygdala (**B**) ROIs. The intensity of negative affect during each condition was represented by the BOLD signal changes of the negative block versus of the neutral block. (**C**) An illustration of amygdala activation among watching, GI and RII condition (FWE corrected p=.01 and an extent of 10 voxels). Error bars = SEM, ns stands for not significant, ** means p≤0.01.

The univariate ANOVAs of subjective experience and the percent BOLD signal changes in the key emotion-generative-regions (amygdala and insula) were conducted among the three conditions (watching, GI and RII). At both the behavioral and neural level, the intensity of negative affect during each condition was represented by the negative emotion index minus the neutral emotion index, and its higher values mean more negatively emotional intensity during the condition.

#### Subjective experience

There was a significant difference across the three conditions (**Figure 2A**), both before (F(2,50)=29.82, p<0.001, η^2^=0.54) and after (F(2,49)=6.62, p<0.01, η^2^=0.22) taking habitual reappraisal as a covariate. Bonferroni post hoc comparisons indicated that the implementation intention condition (M=2.74) had significantly lower negative emotional level than the watching condition (M=3.98, p<0.001) and the GI condition (M=4.11, p<0.001). The watching condition and the GI condition showed no significant differences in negative emotion experience, p=1.

#### Neural responses in emotion-generative ROIs

There were significant main effects in bilateral amygdala responses (left/right amygdala: F(2,50)=6.43/3.62, p=0.003/0.03, η^2^=0.20/0.13; **Figure 3A/B**). Bonferroni post hoc comparisons that in the left amygdala, PSC (M=0.25) was smaller during implementation intention than in watching conditions (M=0.53, p=0.008). The watching and the GI (0.37) condition showed no significant differences, p=0.10. Similarly, in the right amygdala, PSC during implementation intention condition (M=0.20) was also smaller than the watching condition (M=0.36, p=0.02). The watching condition and the GI (M=0.28) condition showed no significant differences, p=0.68. However, no significant main effects were found in the left insula (F(2,50)=0.90, p=0.42, η^2^=0.04), or in the right insula responses (F(2,50)=0.51, p=0.60, η^2^=0.02).

These findings showed that forming implementation intention was effective in realizing emotion-regulatory goals, reducing negative feelings and disgust-related neural activations (bilateral amygdala).

### Cognitive Costs during Implementation Intention: behavioral and ROI analyses

The univariate ANOVA analysis of both subjective perceived effort and the percent BOLD signal changes in the bilateral dlPFC and dACC among the three conditions were conducted.

#### Subjective Report

No significant differences concerning the subjective effort in negative emotion regulation emerged among the watching (M = 2.57), the GI (M = 2.69) and the implementation intention conditions (M = 2.27, F(2,50)=1.82, p=0.17, **Figure 2A**). The selfreported difficulties in coping negative emotions were significantly different among the watching (M = 2.42), GI (M = 2.35) and RII conditions (M = 1.73, F(2,50)=7.64, p<0.01, η^2^=0.23, **Figure 2A**). The RII condition was linked with a significantly lower report of difficulties than the watching (F(1,25)=14.46, p<0.01, η^2^=0.37) and GI conditions (F(1,25)=7.66, p=0.01, η^2^=0.23), whereas the latter two conditions showed no significant differences (F(1,25)=0.19, p=0.66).

#### Neural responses in cognitive-control-related ROIs

To check whether implementation intention engenders voluntary control in the neural level, we directly tested the BOLD signal changes of key nodes of the frontoparietal control network, including bilateral dlPFC and dACC. The main effects of BOLD responses in all three regions were not significant (all p>0.10) (**Figure 2B**).

### Whole brain analyses

Moreover, to test whether, in addition to the regions of interest, other regions were affected by RII, a 3-by-2 repeated-measures ANOVA was run in a whole-brain analysis with picture valence and type of strategies as factors. The strongest interaction effect was found in the left mPFC, vmPFC and postcentral regions (**Figure 4** and Table 2). Follow-up t tests showed that the difference between negative and neutral block was significant during both GI (left mPFC: t=4.63, p<0.001; vmPFC: t=2.58, p<0.02) and watching conditions (left mPFC: t=6.05, p<0.001; vmPFC: t=8.85, p<0.001) but not during RII (left mPFC: t= −1.86, p=0.075; vmPFC: t=1.18, p=0.25) in left mPFC and vmPFC. The difference between negative and neutral block at the postcentral region was only significant during watching condition (t=7.22, p<0.001) but not during GI (t=1.28, p=0.21) and RII (t=0.14, p=0.89) (**Figure 4**). These findings of whole-brain-analysis are consistent with those of ROI-analysis, indicating that RII did not increase disgust-related neural processing in the prefrontal regions.

**Figure 4.**
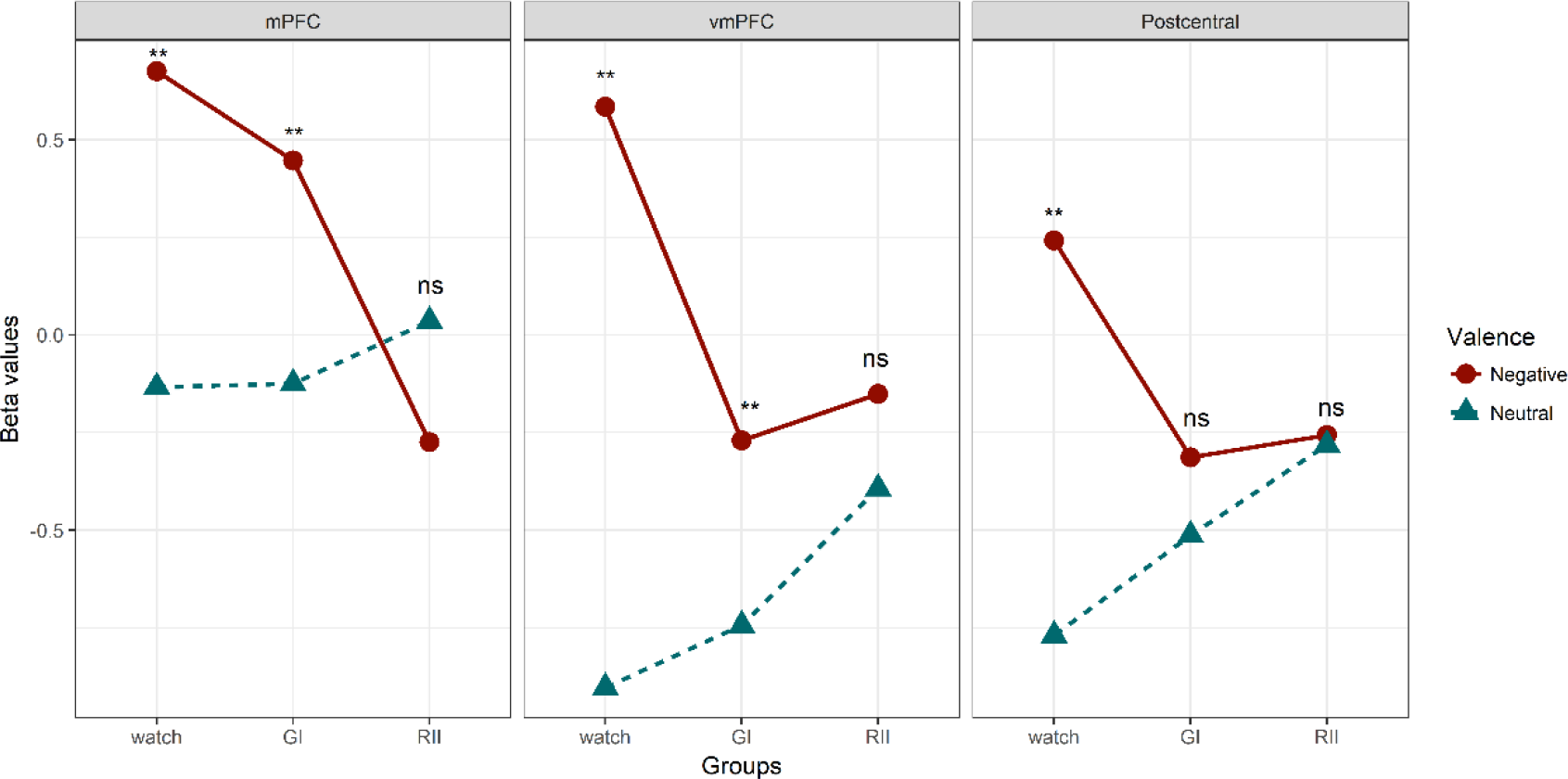
Changes of beta values of activation clusters for the interaction effects (valence*strategy). ** means p≤0.01, ns stands for not significant.

**Table 2.**
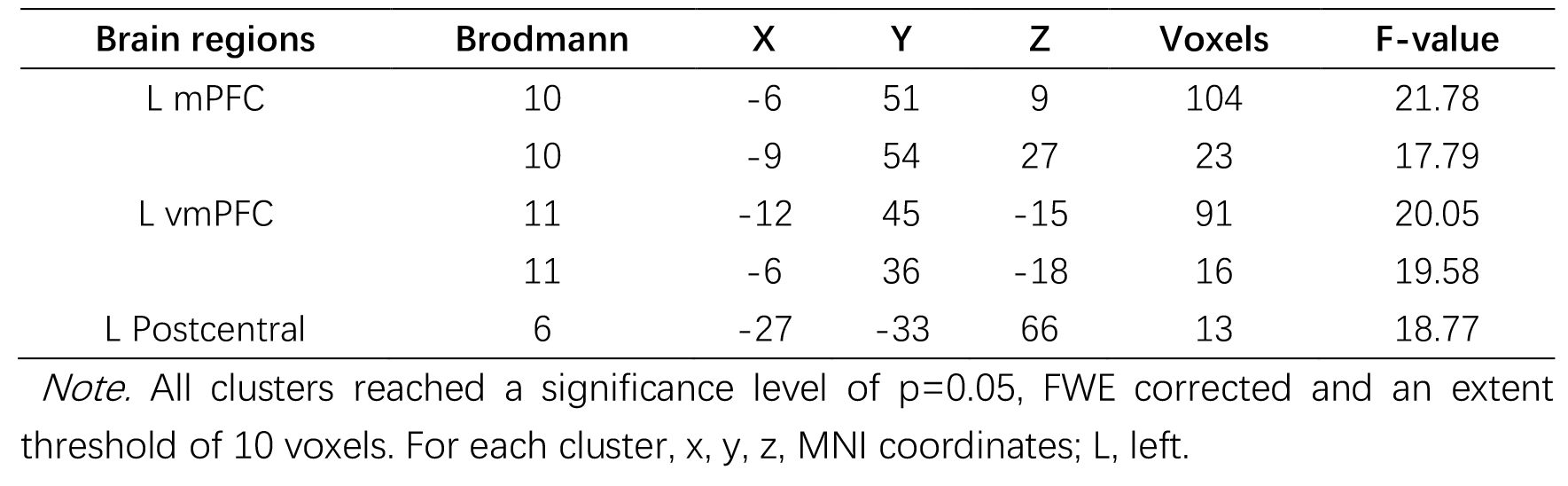
Voxel-wise Group Activations by 3-by-2 ANOVA

Together, these behavioral and neuroimaging data showed that RII facilitated emotion regulation of disgust without costing more cognitive resources compared to watching and GI conditions.

### Task-related FC analyses

We applied task-related FC analysis to the 3-by-2 experimental datasets by considering ROIs as nodes and the block-by-block FC between each pair of ROIs as edge strength. After computing the ROIs pair-wise correlation matrix of each condition for each participant, we conducted a 3-by-2 repeated-measures ANOVA of FC with picture valence and type of strategies as factors at the group level.

The strongest interaction effect was found in 12 pairs of ROI-to-ROI FC (corrected for multiple comparisons via FDR) (see **S2 Table**). Planned comparisons for each FC were then conducted by testing how the FC intensity difference between negative and neutral blocks varies across the regulation conditions. Specifically, we were mainly interested in the contrasts GI and RII versus watching condition and RII versus GI. According to the principles of cognitive subtraction, the contrasts GI and RII versus watching condition should reflect the FC underlying the intentional and automatic pursuit of emotion regulation goals (GI and RII), respectively. The contrast RII versus GI should reflect FC related to the differences of automation between the goal-directed GI and stimulus-driven RII. The results of contrasts GI and RII versus watching condition showed close similarities: four significant FCs for the contrast GI versus watching condition and two of these four FCs for the contrast RII versus watching condition (see **Table 3** and **Figure 5**). Further, seven FCs showed significant decreases in FC intensity during RII than GI (see **Table 3** and **Figure 5C**). These seven FCs, as discussed later, may constitute an interactive neural system underpinning online emotion-related coping. Accordingly, the reduced functional coupling in this system suggests less mobilization of online processing resources for the attainment of emotion regulatory goal during RII.

**Table 3.** Planned Comparisions of Functional Connectivity Strength

| FC | t-value |
| --- | --- |
| **GI vs Watching** |  |
| L vACC - R Precuneus | 4.78 |
| L vACC - R SMG | 4.77 |
| L vACC - L Insula | 4.67 |
| R ITG- L MTG | 2.86 |
| **RII vs Watching** |  |
| L vACC - L Insula | 2.82 |
| L vACC - R Precuneus | 2.53 |
| **RII vs GI** |  |
| R Lingual Gyri - R Putamen | -6.22 |
| R ITG - L MTG | -6.13 |
| R Paracentral Lobule - R STG | -4.66 |
| R Putamen - L Rolandic Operculum | -5.98 |
| R Postcentral Gyri - R Paracentral Lobule | -5.01 |
| R IPL- R SPL | -4.70 |
| L vACC - R SMG | -4.49 |

*Note.* All connections reached a significance level of two-tailed $p < 0.05$ . FC, Functional Connectivity. For each connection, L, left; R, right. Inferior Temporal Gyri=ITG, Middle Temporal Gyri=MTG, Superior Temporal Gyri=STG, Inferior Parietal Lobule=IPL, Superior Parietal Lobule=SPL, SMG=Supramarginal Gyri.

Moreover, we conducted a correlation analysis between the FC intensity and the subjective difficulty index during RII relative to GI, to investigate whether the survived FC could predict behavioral marker of cognitive efforts. Both FC and the index of regulatory difficulty were computed by using GI minus RII. We focused on subjective difficulty index because it was related to negative experiences (r=0.39, p=0.024) during GI relative to RII after a correction of FDR 0.05 (**Figure 6A**), while cognitive effort index was not (r=0.33, p=0.049).

The correlation analysis demonstrated that three of seven FC intensity were positively correlated with subjective difficulty: R Putamen - L Rolandic Operculum, r=0.54, p=0.002; R Lingual Gyri - R Putamen, r=0.35, p=0.04; R Paracentral Lobule - R STG, r=0.41, p=0.02 (**Figure 6BCD**). However, only the correlation of R Putamen - L Rolandic Operculum survived a FDR of 0.05 correction for multiple comparisons.

**Figure 5.**
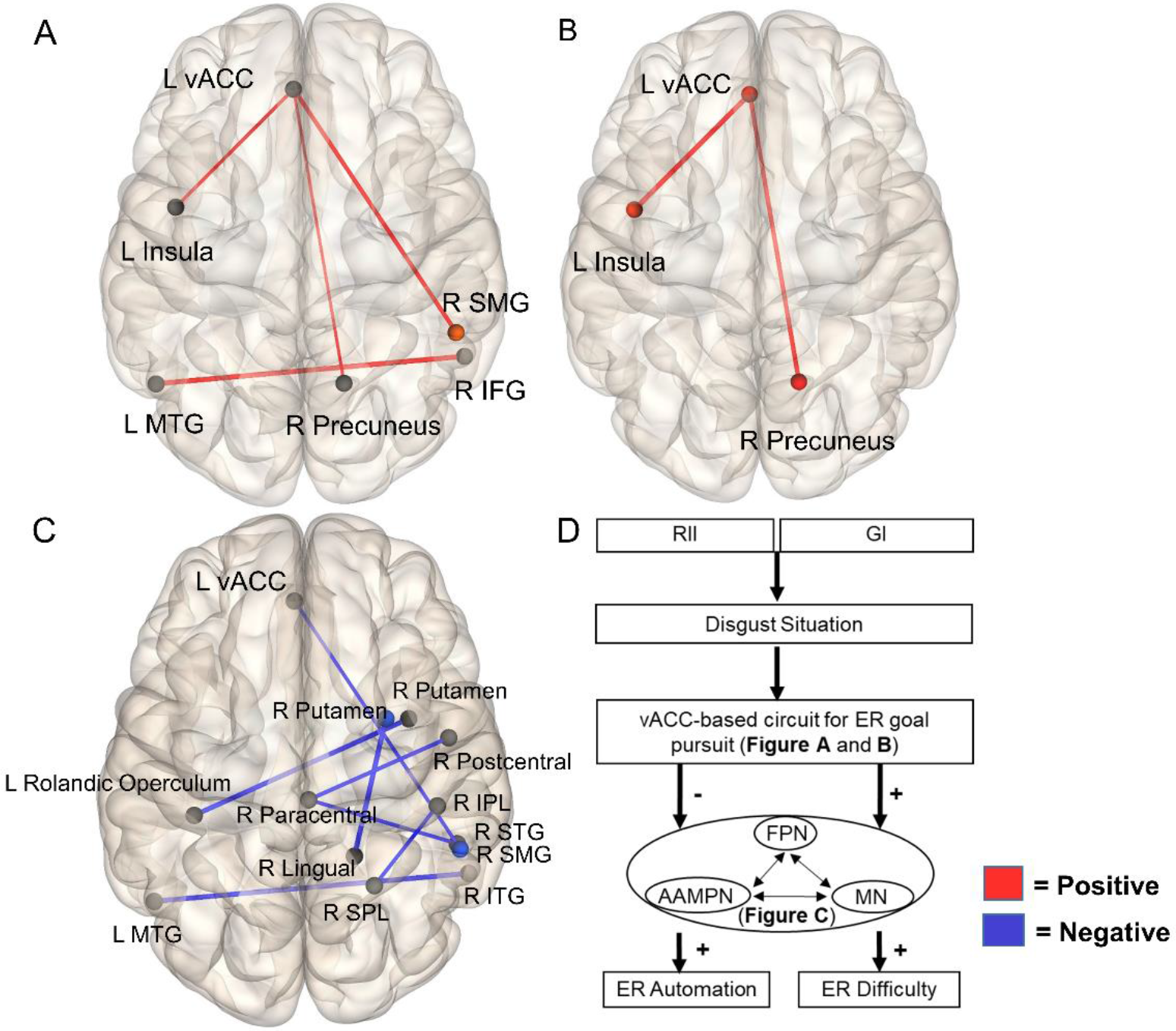
Functional connectivity patterns of the contrasts GI (**A**) and RII (**B**) versus watching condition and the contrast RII versus GI (**C**). The connections (edges) between ROIs marked in red mean that GI or RII relative to watching show greater FC strength, and those marked in blue mean that RII relative to GI shows weaker FC strength. ITG = Inferior Temporal Gyri, MTG = Middle Temporal Gyri, SMG=Supramarginal Gyri. STG = Superior Temporal Gyri, IPL = Inferior Parietal Lobule, SPL = Superior Parietal Lobule, SMG=Supramarginal Gyri. (D) The dynamic architecture of the FC mechanisms underlying GI and RII. FPN, Frontoparietal Network; AAMPN, Aversive Anticipation and Motor Planning Network; MN, Memory Network; ER, Emotion Regulation.

**Figure 6.**
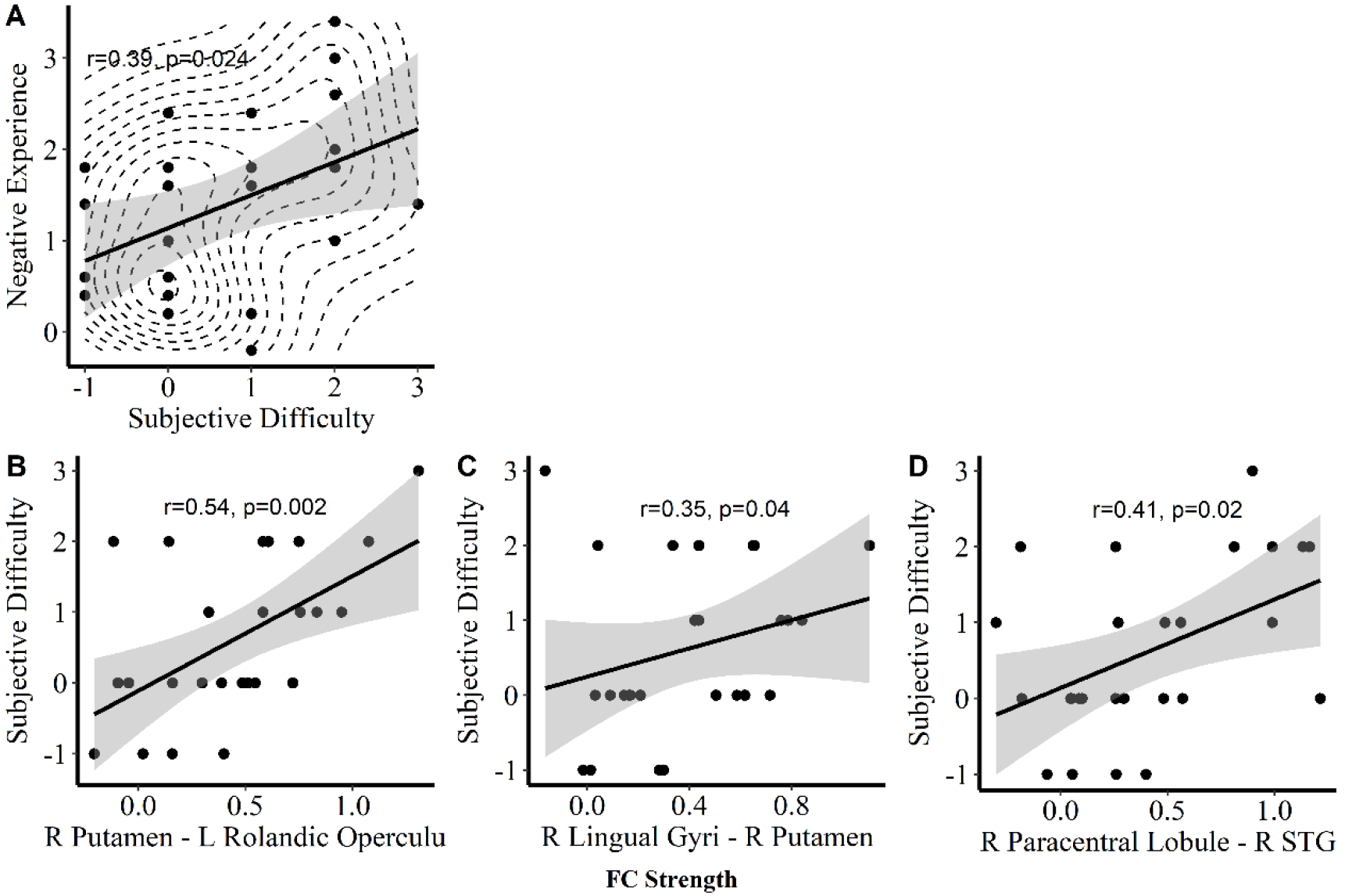
The correlation analysis showed that during GI relative to RII, subjective regulatory difficulty was related to greater negative experiences (**A**), and that the changes of FC intensity were related to greater subjective regulatory difficulty. Results of R Putamen - L Rolandic Operculum (**B**) survived a FDR of 0.05 correction for multiple comparisons, whereas R Lingual gyri – R Putamen (**C**) and R Paracentral Lobule - R STG (**D**) did not. Contour line in **A** was used for dealing with overplotting.

### Discussions

Using behavioral and neuroimaging approaches, this experiment examined the automatic emotion-regulatory effects of RII and their underlying functional connectivity mechanisms. Consistent with previous studies of implementation intention (Eder, 2011; Gallo & Gollwitzer, 2007b; Gallo et al., 2012; Gomez et al., 2015), our results revealed that RII effectively decreased both negative experiences and emotion-related activity in bilateral amygdala. Importantly, these emotion-regulatory effects were not achieved at the cost of greater involvement of cognitive control resources, as the RII did not increase self-reported efforts and control-related prefrontal activations. Furthermore, results of FC analysis demonstrated close similarities between the contrast GI versus watching condition and the contrast RII versus watching condition in vACC-based FCs, and the connectivity strength was decreased during RII than GI in seven distributed FCs.

In comparison with the watching condition, the RII effectively reduced the emotional experiences and activation of bilateral amygdala. By contrast, the GI and watching conditions showed no significant differences at both behavioral and neural indices. These findings coincided with previous findings that forming implementation intentions down-regulated the subjective experience of negative emotions (Gallo & Gollwitzer, 2007a; Gallo et al., 2012), amygdala activation (Hallam et al., 2015) and occipital P1 event-related potential amplitudes (Gallo et al., 2009). Our findings confirmed that RII can regulate disgust effectively at both behavioral and neural level.

Importantly, we collected both behavioral and neuroimaging data of cognitive costs to test the automatic, effortless regulation during RII compared to GI or control condition. On the one hand, we observed no enhancement of subjective efforts but reduced control difficulty during RII compared to the other two conditions. On the other hand, RII did not increase the activity of control-related regions including bilateral dlPFC and dACC compared with watching and GI during ROI analysis. The dlPFC and dACC have been suggested to be generally involved in cognitive-demanding tasks, such as working memory (Owen, McMillan, Laird, & Bullmore, 2005; Wager & Smith, 2003), decision making (Paulus et al., 2001) and voluntary emotion regulation (Goldin et al., 2008). The increased activation of these regions are considered to represent increased cognitive control (Goldin et al., 2008; Wager & Smith, 2003). Moreover, the results of whole brain analysis showed that RII decreased the activation in mPFC and vmPFC in comparison with watching and GI, two regions that also plan important roles in appraisal, expression and regulation of negative emotion, similar to the cognitive control functions of dlPFC and dlPFC described above (Etkin, Egner, & Kalisch, 2011; Motzkin, Philippi, Wolf, Baskaya, & Koenigs, 2015; Schiller & Delgado, 2010; Urry et al., 2006). These behavioral and neuroimaging evidence consistently shows that emotion regulation by RII does not involve cognitive control mechanism to achieve regulatory purposes.

Moreover, FC analysis showed close similarities between the contrast GI versus watching condition and the contrast RII versus watching condition. Specifically, the intensity of FC between left vACC and two nodes (right precuneus and left insula) was increased during both GI and RII than watching conditions. Given that RII builds upon the GI and the context-response association, it is reasonable to infer that these two FCs may be necessary for self-related emotion-regulatory goal pursuit, regardless of degree of task automation. The left insula has been suggested to be a key node of SN (Power et al., 2011), critical for developing and updating motivational states with specific associated actions (i.e., goals) (Kinnison, Padmala, Choi, & Pessoa, 2012; Wager & Barrett, 2004). And the vACC and precuneus are hubs of DMN, involved in self-relevant information processing (Cavanna & Trimble, 2006; Power et al., 2011). Given the close association between SN and DMN (Bonnelle et al., 2012), these two networks may interact to mark the emotionally salient stimuli and then to process it directed by the self-relevant goals (“I will not get disgusted”). Of these three nodes, the vACC was of particular interest, because of its involvement in assessing the salience of emotional and motivational information and the regulation of emotional responses (Bush, Luu, & Posner, 2000; Phillips, Ladouceur, & Drevets, 2008). The vACC is also anatomically connected with emotion-generative regions (e.g. amygdala and anterior insula) (Devinsky, Morrell, & Vogt, 1995; Ghashghaei, Hilgetag, & Barbas, 2007), and its dysfunction is implicated in clinical anxiety or depression (Etkin, Prater, Hoeft, Menon, & Schatzberg, 2010; Kaiser, Andrews-Hanna, Wager, & Pizzagalli, 2015).

Beyond the similarities between GI and RII, the connectivity strength was decreased during RII than GI in seven distributed FCs. Previous studies have pointed out that GI is a goal-directed process, whereas RII is a stimulus-driven one (Gollwitzer & Brandstätter, 1997; Webb, et al., 2012). Therefore, we suggest the increased FC strength during GI compared to RII may reflect a goal-directed (top-down) online emotion-related coping that underlies the gap between emotion-regulatory goals and emotion-regulatory success. These seven FCs can be summarized into three networks that may cooperate to perform this process. Firstly, the connection IPL-SPL, as a part of FPN (Corbetta, 1998; Ptak, 2012), is involved in preparing and applying goal-directed (top-down) selection for stimuli and responses (Corbetta & Shulman, 2002), and its activity increases with higher cognitive demand (Seminowicz & Davis, 2007). Secondly, the Putamen-Rolandic Operculum, vACC- SMG, Postcentral-Paracentral Lobule and Paracentral Lobule-STG connections are implicated in aversive anticipation (Postcentral-Paracentral Lobule and Paracentral-STG) and emotion-related motor planning and preparation (Putamen-Rolandic Operculum and vACC-SMG). Specifically, anticipatory activity in right postcentral gyrus and STG is associated with greater emotional responses and decreased regulation success (Denny, Ochsner, Weber, & Wager, 2014), and neural patterns of postcentral gyrus and paracentral lobule are closely related to level of averseness (Sarkheil, Goebel, Schneider, & Mathiak, 2013). Further, SMG is involved in planning of goal-oriented actions (Tunik, Lo, & Adamovich, 2008). Putamen and rolandic operculum also play similar roles in motor planning (Alexander & Crutcher, 1990; DeLong et al., 1984) and execution (Brown et al., 2009; Ciccarelli et al., 2005), respectively. Lastly, ITG, MTG and lingual gyri, are key nodes of memory systems, mediating interconnected memory functions, like establishment of representations in long-term memory (Rolls, 2000; Squire & Zola-Morgan, 1991) and memory retrieval (Cho et al., 2012).

Together, these three networks may constitute a neural system subserving the goal-directed, online emotion coping mechanism, including components of cognitive control, memory reference and retrieval, as well as aversive anticipation and motor planning. Without antecedent formation of a situation-response association (e.g. if-then plan), participants may have mobilized this system upon receipt of stimulus to achieve their emotion-regulatory goals, leading to higher experienced regulatory difficulty during GI than RII (see **Figure 5D**). Additionally, there are positive correlations between the two FCs (R Putamen- L Rolandic Operculum and Paracentral Lobule-STG) and self-rating of emotion coping difficulty. These results suggest that increased regulatory difficulty is coupled by the higher online mobilization of emotion-related coping network, which provides a possible explanation for the little emotion regulation effect during goal intention.

On the other hand, the decreased FC strength during RII may reflect a greater degree of goal-dependent automaticity (Bargh, Schwader, Hailey, Dyer, & Boothby, 2012), which may result from the if-then plan component of RII, which builds up conditioning of goal-directed responses to anticipated situational cues, and thus automate these responses when the situational cue is encountered (Gallo & Gollwitzer, 2007b; Gollwitzer, 1999). In this regard, our findings are consistent with previous findings that task automatization is accompanied by decreased activation of putamen (Poldrack et al., 2005) and FPN (Mohr et al., 2016). We therefore posit that the FC strength in the emotion-coping network may serve as a neural marker that negatively predicts the automation of emotion regulation. Previous neural mechanism models, influenced by the theory of cortical localization of function, hold that the voluntary and automatic subprocesses of emotion regulation are performed by different neural systems (Gyurak, Gross, & Etkin, 2011; Phillips et al., 2008). And neural regions underpinning cognitive control, such as the lateral prefrontal cortical regions (e.g., dlPFC), are involved in voluntary emotion regulation (Goldin et al., 2008; K. McRae et al., 2012). However, it is unclear to date what neural mechanisms underlie automatic emotion regulation. The current findings, from the perspective of functional integration, posit that the strength of FCs subserving emotional coping may be an important indicator for assessing the automation during emotion regulation.

## Experiment 2

### PURPOSE AND RATIONALE

Because the three emotion regulation conditions (watch, GI and RII) are presented in sequential runs, potential emotional habituation or repeated formation of GI may influence the findings of Experimental 1. Thus, any changes in emotion-generative or cognitive control-related regions may simply reflect habituation or practice effects. To test this possibility, Experiment 2 was conducted to repeatedly present stimuli for watching and GI, two conditions exhibiting little emotion regulation effect, for three times.

Accordingly, we conducted a 2 × 3 ×2 factorial experiment with the between factor selfregulation condition (watching and GI) and the within-factors type of pictures (neutral and disgust) and repetition times (1-3 times). Our previous study observed no significant habituation to repeatedly presented negative stimuli in emotional rating or brain potentials, suggesting that the humans ‘emotional reaction to negative stimuli are resistant to habituation (Long, Yang, Lou, & Yuan, 2015). Also, the AREA (attend, react, explain, adapt) model of affective adaptation holds that the affective reactions to negative events would not decrease significantly until the negative events are fully understood (Wilson & Gilbert, 2008). However, the cues for effective explanation were unavailable during watching and GI. Based on these studies, we predict that repetition of emotional stimuli for watching or GI condition would not decrease one’s emotional responses to disgusting stimuli.

#### Methods

##### Participants

Forty healthy right-handed females completed the study (mean age=20.0), and were randomly assigned for watching and GI groups. All participants had normal or corrected-to-normal vision, reported no history of neurological or psychiatric disorders. Trait anxiety, state anxiety and depression assessment did not differ between the experimental groups, as shown by their scores in the State (STA; t(38)=−1.26, p=0.22) and Trait (TAI; t(38)=0.94, p=0.35) Anxiety Inventory (STAI) (Spielberger, 1970) and Beck Depression Inventory-II (BDI-II, t=0.67, p=0.51) (Beck, Steer, & Brown, 1996). Before admission to the study, all participants gave their written informed consent. This experiment was approved by the Institutional Review Board of the Southwest University and was in accordance with the latest revision of the Declaration of Helsinki.

##### Stimuli

The neutral and disgust pictures used in this experiment were identical to those used in Experiment 1.

##### Procedure

The general procedure was identical to the procedure of Experiment 1 except for the following changes: 1) the Self-Assessment Manikin (SAM) (Bradley & Lang, 1994) were used to assess the valence ratings with respect to each of the block presented, and the valence rating ranged from 1 (very happy) to 9 (very unhappy); 2) the watching or GI tasks needs to be performed for three times.

##### Results and Discussion

The results of three-factorial ANOVAs yielded neither significant interaction effects between repetition times and self-regulation conditions (F(2,76)=1.92, p=0.15, η^2^=0.05), nor the main effects of repetition times (F(2,76)=2.09, p=0.13, η^2^=0.05). There was a significant main effect of picture type (F(1,38)=294.98, p<0.001, η^2^=0.87) (**Figure 7**).

**Figure 7.**
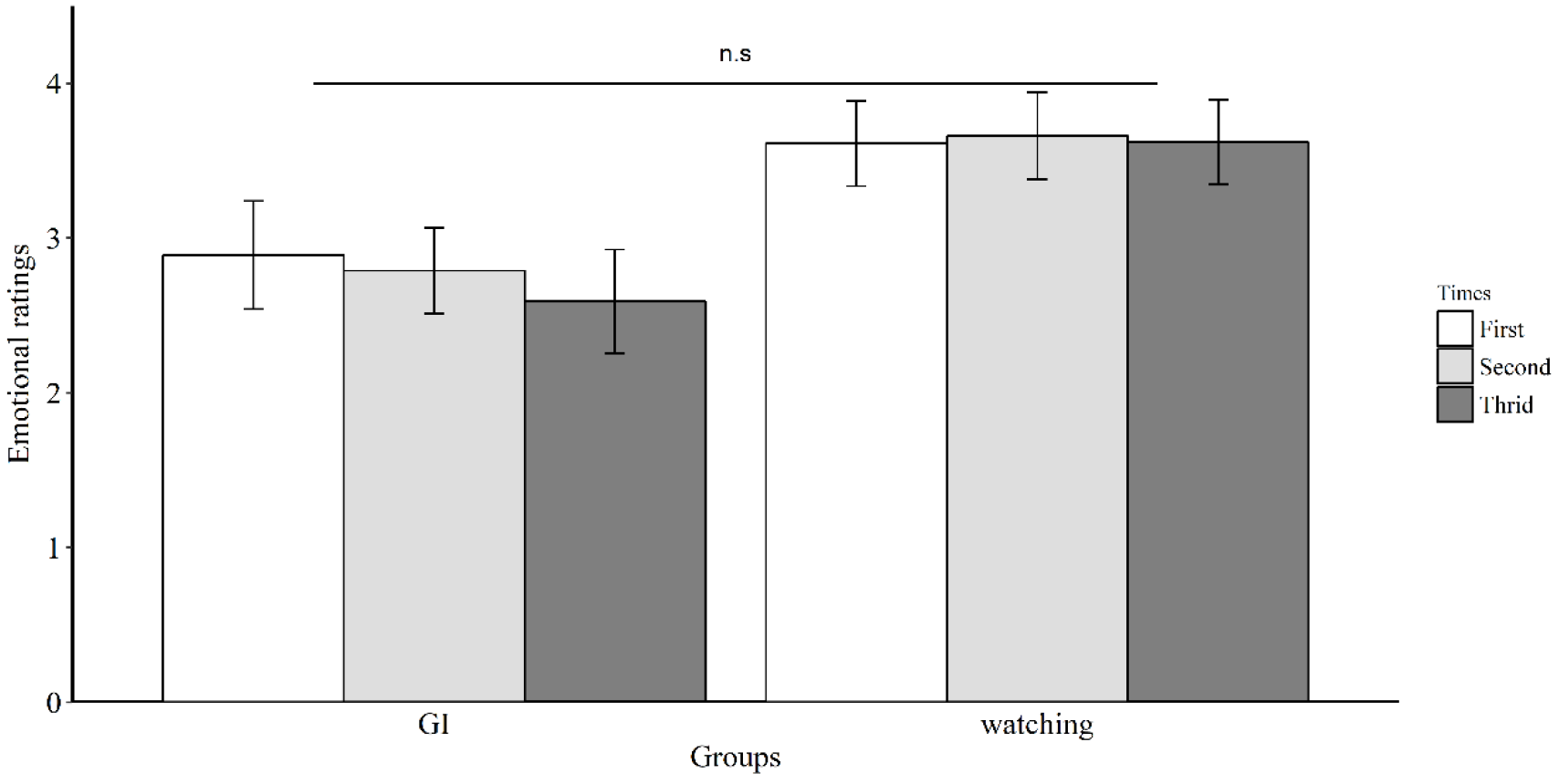
Mean subjective ratings of negative emotions for the watching and GI groups during three times. Error bars=SEM, ns stands for not significant.

These findings are consistent with our prediction, suggesting that emotion rating for these disgust stimuli would not decrease as a result of emotional habituation or GI practice. Importantly, the findings of Experiment 1 and 2 can be explained using the AREA (Attend, React, Explain and Adapt) model of affective adaptation (Wilson & Gilbert, 2008). In this model, Wilson and Gilbert (2008) proposed that people engage in the sequential process of attending, reacting, explaining, and ultimately adapting to affective events. The most novel part of AREA model is that explanation leads to affective adaptation. That is, affective reactions to negative events would not decrease significantly over time until individuals explain negative events successfully. RII, as a combination of cognitive reappraisal and implementation intention, provided participants with effective explanations of disgust stimuli, whereas GI and watching conditions did not. Therefore, the decrease in subjective ratings and amygdala activity during RII compared to watching or GI in Experiment 1 can result exclusively from the regulatory effects of RII.

##### General conclusions and limitations

In summary, we examined whether RII regulates emotion automatically at both the behavioral and neural level and then investigated the neural mechanisms associated with the automatic regulatory effects. We confirmed that RII significantly decreased negative feelings and neural activity in emotion-generative regions without engendering cognitive efforts, as evidenced by the similar or even reduced self-reported efforts and cognitive control-related prefrontal engagement during RII compared to watching and GI. These regulation effects should not be explained by emotional habituation, as Experiment 2 did not show significant emotional habituation for the repetition of the same stimuli. Moreover, RII and GI produced similar FC of vACC to left insula and right precuneus, corresponding to the common goal pursuit component of both strategies. Furthermore, RII relative to GI exhibited weaker FC in distributed brain networks subserving effortful control (e.g. IPL-SPL FC), memory retrieval (e.g. inferior-middle temporal lobes and lingual-putamen FC), aversive anticipation and emotion-related motor planning (e.g. Paracentral-STG, putamen-operculum and vACC-supramarginal gyrus FCs). These findings suggest that RII downregulates disgust automatically, most likely by reducing the online mobilization of neural systems subserving emotion-related coping.

Several limitations need to be acknowledged. First, only healthy participants were studied, and it is therefore unclear whether our results are generalizable to clinical samples. Given the cognitive control deficits in individuals with emotional disorders (e.g., anxiety and depression), implementation intention, as a way of automatic emotion regulation, may facilitate clinical population compared to voluntary strategies. Second, this study only combined implementation intention with cognitive reappraisal. However, there are other emotion regulation strategies, like attentional deployment and expressive suppression (Webb, Miles, & Sheeran, 2012). The neural mechanisms underlying different emotion regulation strategies by implementation intentions may be different. Third, this study focused on the outcome of emotion regulation by implementation intention during aversive stimulation, leaving neural substrates mediating GI and RII formation unmeasured. It is possible that participants paid cognitive resources during formation of implementation intentions before stimulus presentation. Thus, future studies need to assess both formation and application of implementation intention concerning control-related neural underpinnings.

## Acknowledgments

This work was supported by the National Natural Science Foundation of China (grant numbers 31371042, 31400906).

